# xNNPCD identifies regulators of programmed cell death by integrating perturbation transcriptomes with cancer dependency profiles

**DOI:** 10.64898/2026.05.10.724150

**Authors:** Qingyang Yin, Liang Chen

**Affiliations:** Department of Quantitative and Computational Biology, University of Southern California, Los Angeles, CA 90089, United States

## Abstract

Programmed cell death (PCD) encompasses multiple regulated processes whose dysregulation shapes cancer fitness, yet current computational studies largely use known PCD genes for prognosis rather than discovering regulators. We developed xNNPCD, an interpretable neural-network framework that links CRISPR-Cas9 perturbation signatures from CMap to gene dependency profiles from DepMap. The model constrains hidden neurons to five PCD pathways and iteratively refines a prior gene-pathway mask matrix derived from GO, KEGG, and Reactome using pathway-neuron ablation. This converts binary gene-pathway relationships into continuous-valued associations and improves dependency prediction over random forests, standard fully connected multi-layer perceptron, and its own non-iterative variant. The learned matrix recovers annotated death regulators and nominates candidate regulators, including *RPL23A, HSPA5, SNRPA1, SLC6A2*, and *ASAH1*; combined with dependency scores, it further separates pathway coupling from regulatory direction. Transferring the refined relationship matrix and learned weights to compound-induced perturbation data enables *in silico* drug screening, identifying BRD-K19103580 and decitabine as targeted therapeutic agents for apoptosis and ferroptosis, respectively. The pathway-resolved drug profiles can facilitate the rational design of combination therapies targeting complementary PCD pathways to overcome single-pathway resistance. Overall, xNNPCD offers a generalizable, interpretable approach for mapping the regulatory landscape and elucidating the molecular processes of PCD in cancer.

**GRAPHICAL ABSTRACT:** 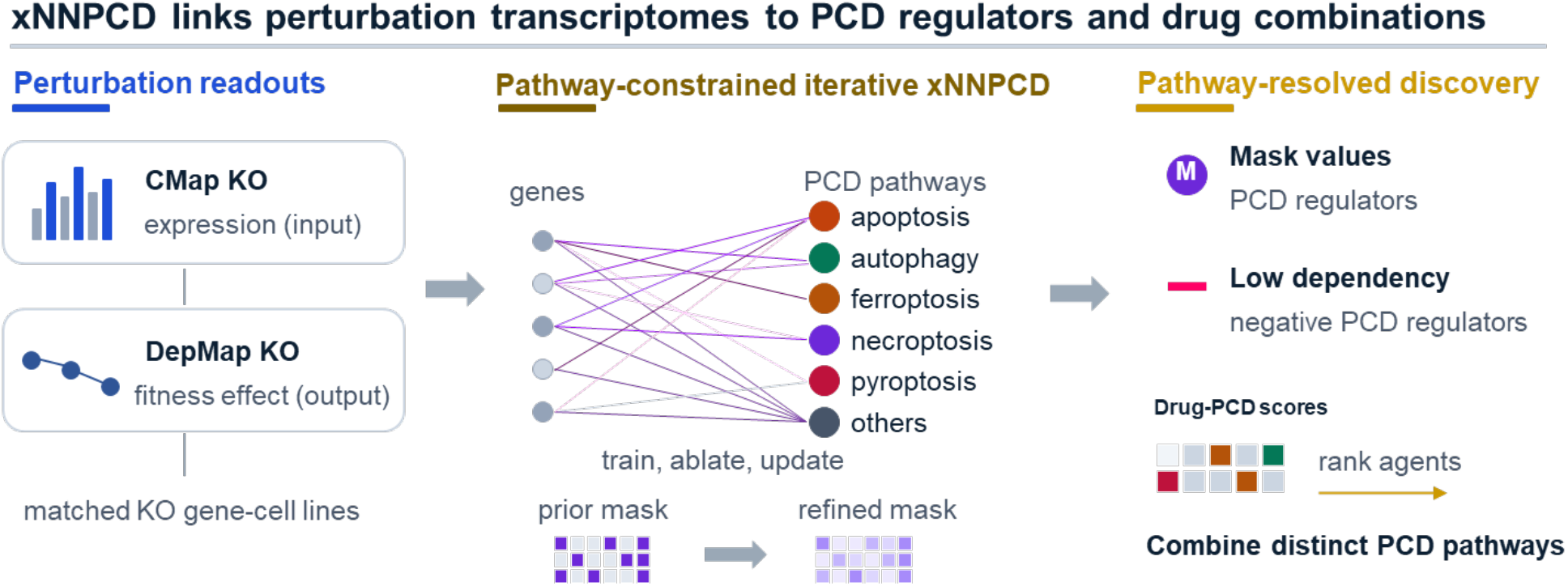

## INTRODUCTION

Programmed cell death (PCD) refers to a collection of genetically regulated mechanisms that eliminate damaged, superfluous, or potentially dangerous cells (1). These mechanisms are indispensable for development, tissue homeostasis, and immune defense (1-3). Multiple molecularly distinct forms of PCD have been characterized, from classical apoptosis and autophagy to more recently defined modalities such as ferroptosis, necroptosis, and pyroptosis, each governed by dedicated signal cascades (4,5). Crucially, extensive crosstalk among these pathways establishes a robust cell-death network in which the activation of one modality can potentiate or suppress another (6). Within this network, regulators that restrain death execution, often termed negative regulators, are critical for maintaining cell survival; their loss can unleash uncontrolled cell death, while their overexpression can confer pathological survival advantages that contribute to disease (7).

Computational approaches to PCD have advanced along two distinct tracks. On one hand, mechanistic modeling using ordinary differential equations, Boolean networks, Petri nets, and cellular automata has been the standard framework for simulating death-pathway dynamics (8). On the other hand, recent data-driven studies have used predefined PCD gene sets to build cancer prognostic signatures, linking PCD-associated expression programs to survival, immune state, immunotherapy response, and drug sensitivity (9-11). While neural network-based models have recently emerged, their use has been largely restricted to image-based morphological classification of dying cells (12,13). Thus, existing data-driven approaches support the clinical relevance of PCD programs, but they do not identify new regulators or assign genes to specific PCD pathways.

Two large-scale functional genomics resources make it possible to study PCD regulation from complementary perturbation readouts. The Connectivity Map (CMap), generated within the LINCS project, catalogs genome-wide differential expression signatures elicited by diverse perturbations, including CRISPR-Cas9 gene knockouts and small-molecule treatments (14,15). The Cancer Dependency Map (DepMap) measures the fitness consequence of each gene loss across cancer cell lines, reported as dependency scores that serve as an inverse proxy for gene essentiality and enable broad applications in cancer genomics (16-18). For shared knockout-cell line pairs, these datasets provide both the molecular response to losing a gene and the survival effect of that loss. This pairing is well-suited for studying PCD regulation because the expression signature can reveal which death-pathway is perturbed, whereas the dependency score helps indicate whether loss of the gene compromises cell fitness. Knockout of a positive regulator may weaken a death-pathway transcriptional program, while knockout of a negative regulator may release or activate it; in both cases, the expression signature can reveal functional coupling to a PCD pathway. Dependency scores then help distinguish regulatory direction: loss of a negative regulator is expected to reduce fitness and produce a strongly negative dependency score, whereas loss of a positive regulator is expected to have a weaker or more context-dependent fitness effect.

To translate these paired perturbation readouts into pathway-level regulator discovery, we developed xNNPCD, an interpretable neural network framework that predicts cell fitness effects while preserving PCD pathway structure. xNNPCD takes CMap knockout-induced expression signatures as input and predicts DepMap dependency scores as output. Unlike fully connected neural networks, which can capture nonlinear associations but do not reveal which biological pathway drives a prediction, xNNPCD uses a pathway-constrained hidden layer: five neurons correspond to apoptosis, autophagy, ferroptosis, necroptosis, and pyroptosis, as well as an additional “others” neuron captures non-PCD effects on cell fitness. Input genes are connected to pathway neurons through a gene-PCD mask matrix, where each entry represents the strength of association between a gene and a PCD pathway. The mask is initialized from prior gene-PCD annotations, so the starting network topology reflects curated biological knowledge. In this design, learned mask values quantify how strongly each gene is coupled to each PCD pathway, while dependency scores provide complementary information about whether loss of that gene reduces cell fitness.

xNNPCD further refines this initial gene-PCD mask through an iterative update procedure. The gene-PCD relationship matrix is initialized from GO, KEGG, and Reactome annotations, and then the model is trained to predict dependency scores from knockout-induced expression signatures. After each training round, pathway-neuron ablation estimates how much each PCD pathway contributes to the predicted fitness effect of each knockout. These pathway-specific importance scores are then used to update the initial binary mask into a continuous-valued matrix, allowing xNNPCD to recover both known and previously unannotated gene-PCD associations. The refined matrix identifies regulators by scoring each gene’s association with specific PCD pathways, while more negative dependency scores help prioritize negative regulators whose loss compromises cell fitness. Finally, transferring the refined mask matrix and learned weights to compound-induced perturbation data enables pathway-specific drug screening, providing a foundation for designing combination regimens in which drugs targeting distinct PCD pathways are paired to overcome resistance to any single death modality.

## MATERIAL AND METHODS

### Input datasets

#### The CMap dataset

We utilized the Connectivity Map (CMap) dataset from the Library of Integrated Network-based Cellular Signatures (LINCS) project (CMap LINCS 2020) (14). Expression was measured using the L1000 high-throughput assay, which directly profiles 978 landmark genes and computationally infers the expression of 11,350 additional genes.

For the primary xNNPCD model, we used the Level 5 data from CRISPR-Cas9 gene knockout experiments. Each Level 5 signature is a moderated z-score (MODZ) vector that quantifies the differential expression profile relative to population controls (19). A signature corresponds to a specific gene knockout in a given cell line, yielding a total of 142,901 signatures covering 5,167 unique target genes across 27 cell lines. Because the dataset retains technical replicates (e.g., from distinct detection wells), multiple signatures may exist for the same gene-cell line pair.

We additionally used the Level 5 compound-treatment signatures, initially containing 720,216 samples. For each compound-cell line combination, we retained the single experiment with the longest perturbation time (typically 24 hours) and the highest dose (typically 10 µM), yielding 167,765 signatures.

#### The DepMap database

We downloaded the CRISPR-Cas9 screening dataset from the DepMap portal (24Q2 release), a cell-line-by-gene dependency matrix encompassing 17,931 genes across 1,095 cell lines (16,17). Each dependency score is derived from the log-fold changes in sgRNA read counts between pre- and post-knockout time points and serves as an inverse proxy for gene essentiality: a more negative score indicates that the cell line is more dependent on that gene for survival, and thus the gene is more essential.

To pair phenotypic and transcriptomic data, we intersect DepMap and CMap by matching common cancer cell lines and knocked-out genes, yielding 115,472 samples with paired differential expression signatures and dependency scores. For gene-cell line pairs represented by multiple CMap replicates, all replicates were assigned the same dependency score.

#### The CTRP database

We obtained drug sensitivity data from the Cancer Therapeutics Response Portal (CTRP v2.0) (20), using the area under the dose-response curve (AUC) as the sensitivity metric. AUC values were normalized by dividing each value by the maximum AUC observed for that compound across all cell lines. After averaging normalized AUCs over replicate experiments, we matched drug-cell line pairs with the filtered CMap compound signatures, yielding a final cohort of 6,665 samples.

#### Data splitting

Gene identifiers in the CMap dataset were mapped from Entrez IDs to gene symbols using the mygene Python library (21) to ensure consistency with pathway annotations. Both the CRISPR-Cas9 knockout and compound datasets were randomly partitioned into training (80%), validation (10%), and test (10%) sets using a fixed random seed.

### Gene-PCD pathway relationship matrix

We constructed a binary gene-PCD pathway relationship matrix *M*^(*orig*)^ by aggregating annotations from GO (22), KEGG (23), and Reactome (24) for five PCD pathways: apoptosis (GO:0006915, hsa04210, R-HSA-109581), autophagy (GO:0048102, hsa04140), ferroptosis (GO:0097707, hsa04216), necroptosis (GO:0097300, hsa04217, R-HSA-5213460), and pyroptosis (GO:0141201, R-HSA-5620971). The resulting matrix has dimensions 12,328 × 6, corresponding to all CMap genes and six categories (five PCD pathways plus “others”). An entry 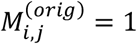 indicates that gene *i* is annotated to pathway *j*; otherwise, it is 0. The “others” category is set to 1 for all genes, providing a fully connected channel to capture fitness effects outside the five defined PCD pathways.

Additionally, to support subsequent downstream analyses, we identified genes associated with the negative and positive regulation of these PCD processes by querying their respective GO terms: apoptosis (GO:0043066, GO:0043065), autophagy (GO:0010507, GO:0010508), ferroptosis (GO:0110076, GO:0160020), necroptosis (GO:0060546, GO:0060545), and pyroptosis (GO:0160028, GO:0140639).

### Neural network structure

The core model is a three-layer neural network: an input layer of 12,328 gene differential expression features, a hidden layer of six neurons (one per PCD pathway plus “others”), and a scalar output layer predicting the dependency score. Connections between the input and hidden layers are governed by the soft mask matrix *M*, initialized as *M*^(*orig*)^ and iteratively refined (see Iterative mask update). The hidden layer activation is computed as *h = f*[(*M*⊙*W*_1_)*x* + *b*_1_], where *x* is the input differential expression vector, *W*_1_ is the weight matrix, *b*_1_ is the bias vector, *f* is the ReLU activation function, and ⊙ denotes the element-wise multiplication (Hadamard product). This operation scales the weights using the soft mask values ranging from 0 to 1, effectively adjusting the strength of each gene-pathway connection. The final dependency prediction *y* is obtained from the linear aggregation of the hidden layer neurons using *y = W*_2_*h* + *b*_2_.

### Ablation analysis

To quantify the contribution of each PCD pathway to a gene’s predicted effect on cell survival, we performed neuron-level ablation. For each knocked-out gene *i*, we extracted its sample subset *S*_*i*_ and obtained the mean baseline prediction 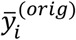 from the intact model. We then set the post-activation value of pathway neuron *p* to zero while keeping all other neurons unchanged, yielding a mean ablated prediction 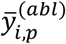. The importance score is defined as

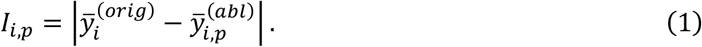

Importance scores were min-max normalized across all genes for each pathway:

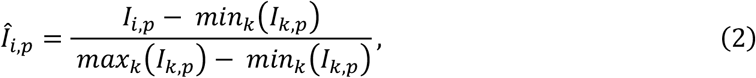

yielding the proposed relationship vector *Î*_*i*,∙_ for gene *i*. The “others” neuron was excluded from ablation; its mask entries remained fixed at 1.

### Iterative mask update

The mask matrix is refined through a two-step update combining biological priors with momentum smoothing. Given the normalized importance matrix *Î* from ablation analysis, an intermediate candidate is computed as:

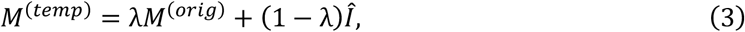

where *λ* ∈ [0,1] controls the weight of the biological prior. The mask for iteration *t* is then updated with momentum:

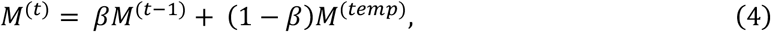

where *β* ∈ [0,1] governs temporal smoothing. At iteration *t =* 0, the mask is initialized as *M*^0^ *= M*^(*orig*)^, network weights were initialized using the Kaiming uniform method (25), and all biases were set to zero. For subsequent iterations (*t* > 0), the model inherited weights and biases from the previous iteration to enable continuous learning. The maximum iteration count and optimal values of *λ* and *β* are determined by hyperparameter tuning.

### Hyperparameter tuning

#### Neural network hyperparameters

We conducted a grid search over learning rate, weight decay, and batch size prior to iterative training. Each configuration was trained on the training set and evaluated on the validation set, with a maximum of 200 epochs and early stopping triggered after 10 consecutive epochs without improvement in validation loss. A step learning rate scheduler decayed the learning rate by a factor of 0.5 every 5 epochs. The optimal configuration was: learning rate = 0.01, weight decay = 0.001, batch size = 128, with minimum validation loss at epoch 43. These settings were fixed for all subsequent experiments.

#### Mask update hyperparameters

We tuned *λ* and *β* via grid search, monitoring validation *R*^2^ at each iteration, with a maximum of 10 iterations and early stopping after 2 consecutive iterations without improvement. The optimal values were *λ =* 0 and *β =* 0.5, with 5 iterations. Five-fold cross-validation on the combined training and validation sets confirmed these settings. The final xNNPCD model was retrained on the combined training and validation data before evaluation on the held-out test set.

### Prediction of drug sensitivity

To evaluate translational potential, we adapted the model to predict CTRP drug sensitivity (normalized AUC) from CMap compound-induced differential expression signatures. We implemented a transfer learning strategy: the three-layer architecture retained the final refined mask matrix *M* and was initialized with the final network weights from the gene knockout task. The model was then fine-tuned on the pharmacological data using the same hyperparameter search procedure, yielding: learning rate = 0.01, weight decay = 0.001, batch size = 128, and optimal epoch = 37. The final model was retrained on the combined training and validation data and evaluated on the held-out test set.

To identify pathway-specific drugs, we computed the mean activation value of each PCD pathway neuron *p* across all samples treated with a given drug *d*, yielding a raw importance score *A*_*d,p*_. These scores were standardized into z-scores *Ã*_*d,p*_ across all drugs for each pathway, serving as the final metric for ranking pathway-specific modulators.

### Baseline models

We benchmarked xNNPCD against Random Forests (RF) and Multi-Layer Perceptron (MLP) baselines, and the non-iterative variant of our model (xNN). RF and MLP were evaluated with two input configurations: all 12,328 CMap genes (“all”) and the subset of genes annotated to at least one PCD pathway (“death”). RF used default parameters. The MLP adopted the same three-layer architecture as xNNPCD but with fully connected layers, isolating the effect of the pathway-constrained mask. xNN uses the same pathway-constrained architecture as xNNPCD, but with the original binary mask *M*^(orig)^ fixed throughout training, isolating the effect of iterative mask refinement. Predictive performance was evaluated using the coefficient of determination (*R*^2^) and mean squared error (MSE).

## RESULTS

### The xNNPCD framework

The overall pipeline of xNNPCD is illustrated in Figure 1. The framework utilizes CMap transcriptome-wide differential expression signatures as input to predict DepMap cell fitness dependency scores. Central to the framework is an iterative refinement process that dynamically updates the gene-programmed cell death (PCD) pathway relationship matrix (*M*), enabling the discovery of gene-pathway associations beyond those cataloged in existing databases.

**Figure 1:**
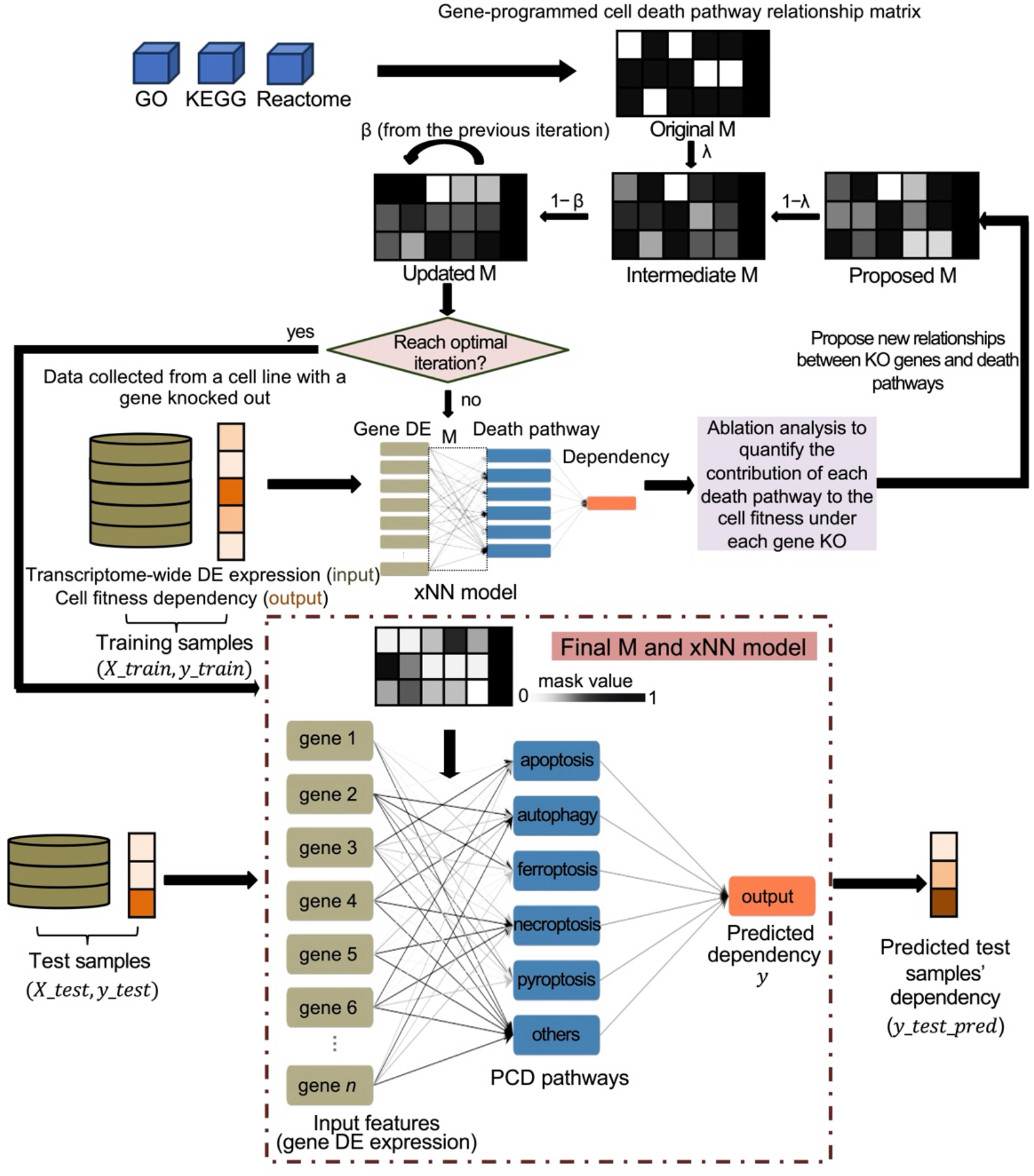
The graphical illustration of the xNNPCD framework. The model takes transcriptome-wide differential expression as input to predict cell fitness dependency, which quantifies gene essentiality for cell survival. The gene-PCD pathway relationship matrix (*M*) is initialized by querying the GO, KEGG, and Reactome databases. In each iteration, the xNN model is trained on the training set. The trained model then undergoes an ablation analysis to quantify the contribution of each cell death pathway to cell fitness under each knocked-out (KO) gene, proposing new relationships between KO genes and death pathways. Subsequently, a composite strategy utilizing convex combination and momentum is applied to update *M*. Upon reaching the optimal iteration, the final *M* and xNN model are deployed on the test set to generate predicted dependencies.

The process begins with *M* initialized as a binary matrix constructed by querying GO, KEGG, and Reactome for established links between genes and five major PCD pathways (apoptosis, autophagy, ferroptosis, necroptosis, and pyroptosis), along with a fully connected “others” channel to capture non-PCD effects on cell fitness. Across the five PCD pathways, a total of 954 unique genes carry at least one annotation, with apoptosis (751 genes) and autophagy (144 genes) being the most represented, followed by necroptosis (147 genes) and ferroptosis (37 genes), and pyroptosis (26 genes) being the least represented. In each iteration, the pathway-constrained neural network (xNN) is trained on the training data, with *M* serving as a soft mask that modulates input-hidden layer connections. The trained model then undergoes ablation analysis (Figure 2): for each knocked-out (KO) gene, we systematically silence each PCD pathway neuron and compare the resulting predictions against the baseline to compute a pathway importance vector. Specifically, the raw importance of pathway *p* for gene *i* is quantified as the absolute difference in mean predictions before and after ablation. After normalization, these importance vectors form a proposed relationship matrix that captures candidate gene-pathway associations. The mask *M* is then updated for the subsequent iteration via a convex combination with the biological prior and momentum smoothing from the previous iteration, ensuring stable convergence. Upon reaching the optimal iteration, the final *M* and trained xNN are used to generate predictions on the held-out test set.

**Figure 2:**
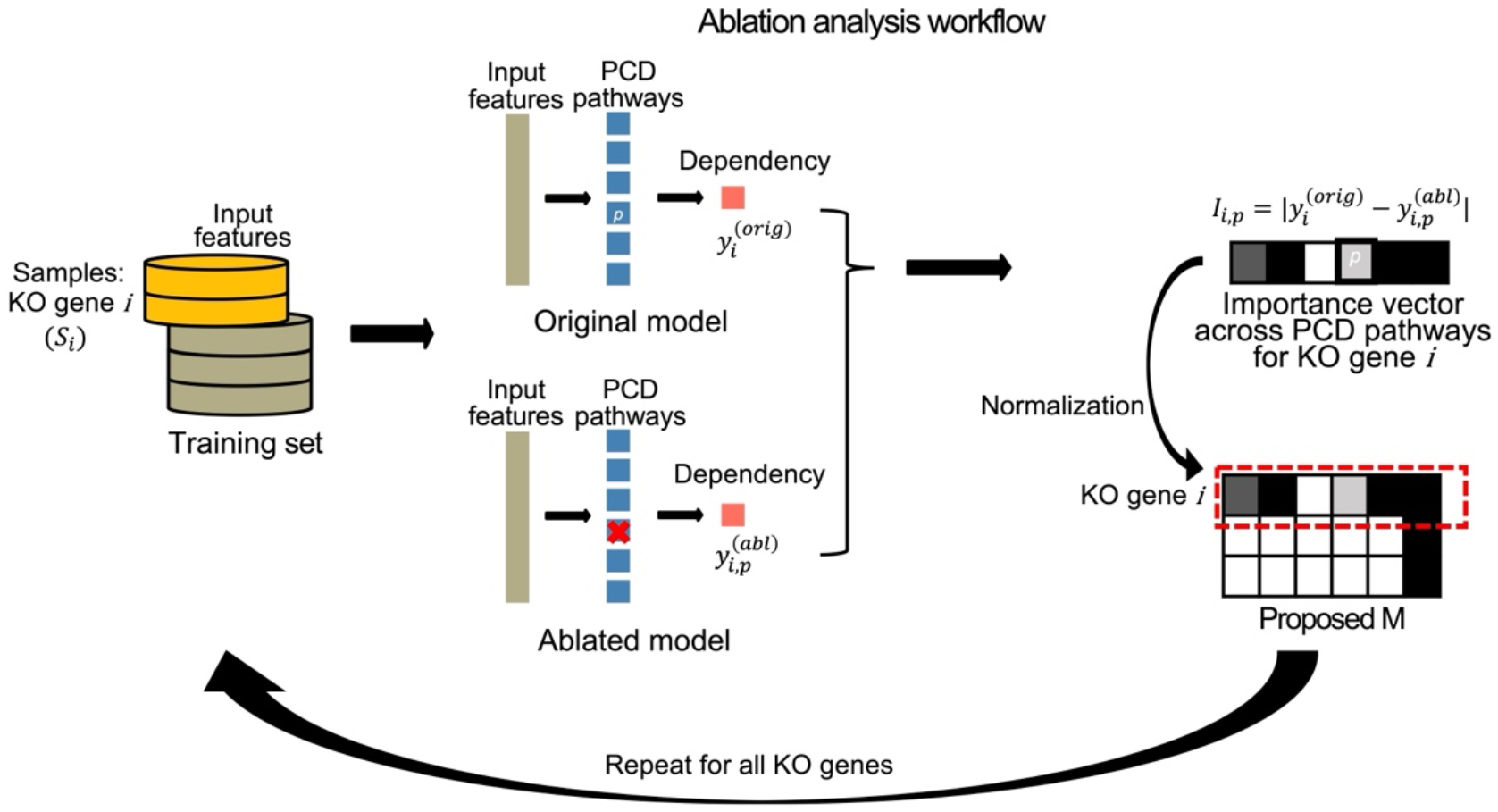
The ablation analysis pipeline. For each knocked-out (KO) gene *i*, the corresponding samples *S*_*i*_ are extracted. Leveraging the weights optimized during the training stage, *S*_*i*_ is first input into the original model to obtain the baseline average prediction 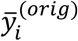. Subsequently, the neuron corresponding to each PCD pathway *p* is ablated, and *S*_*i*_ is re-evaluated to yield a new average prediction 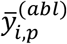. The importance metric is defined as 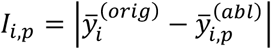. Finally, these scores form the specific row for gene *i* in the importance matrix *I*, which is used to propose new gene-cell death relationships.

### xNNPCD accurately predicts cell fitness dependency

We benchmarked xNNPCD against Random Forests (RF), Multi-Layer Perceptron (MLP), and the non-iterative xNN using CMap differential expression signatures to predict DepMap dependency scores. RF and MLP were each evaluated with two input configurations: all 12,328 CMap genes (“all”) and the 954 known PCD genes (“death”). As illustrated in Figure 3, xNNPCD achieved the highest *R*^2^ score and the lowest MSE across all comparisons.

**Figure 3:**
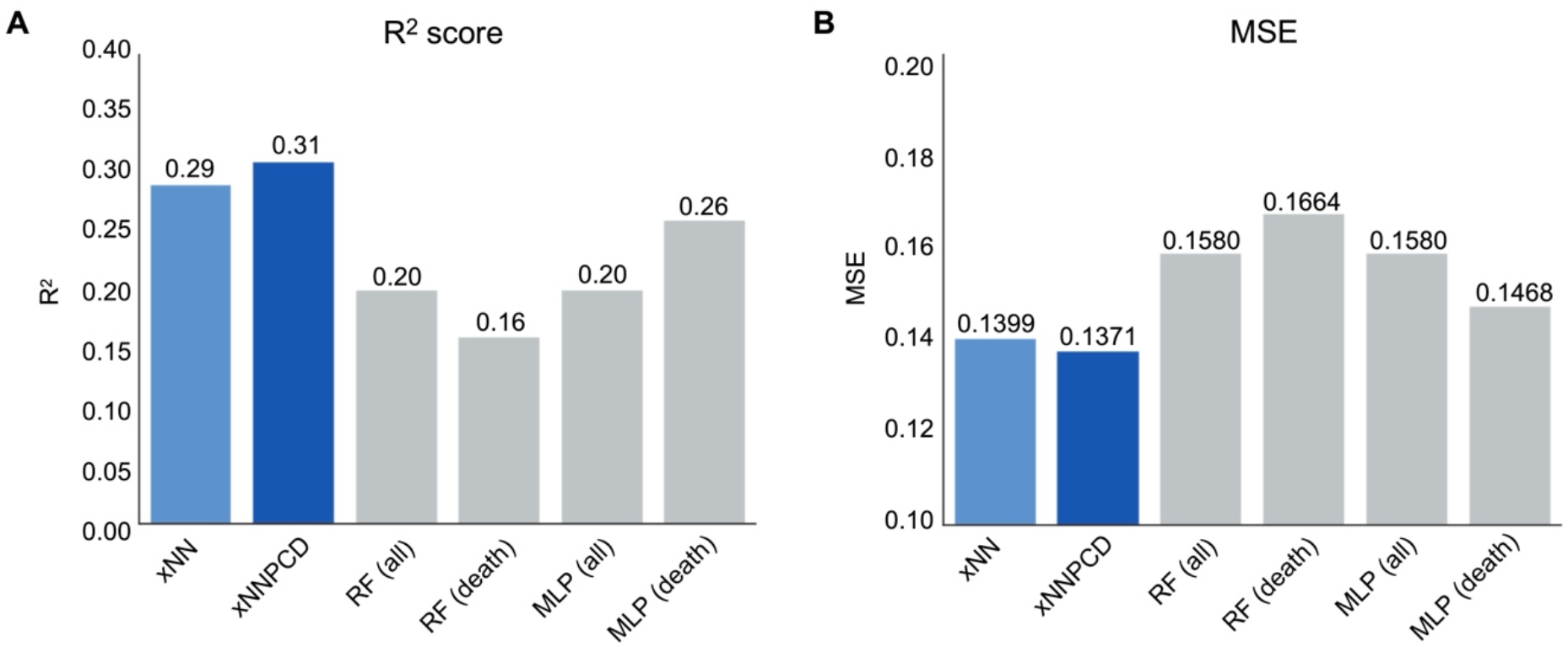
Performance comparison of xNNPCD against baseline methods in predicting cell fitness dependency. The evaluation includes the static xNN (non-iterative), the proposed xNNPCD, Random Forest (RF) models, and Multi-Layer Perceptron (MLP) models. For RF and MLP, two configurations are compared: one utilizing transcriptome-wide features (“all”) and another restricted to known PCD-associated genes (“death”). The predictive performance is quantified by (**A**) the *R*^2^ score and (**B**) the Mean Squared Error (MSE).

The advantage over genome-wide RF and MLP models (“all”) demonstrates the value of encoding PCD pathway structure into the network. Rather than treating all genes as a flat feature vector, xNNPCD groups inputs into pathway-specific neurons, enforcing biologically motivated sparsity that mitigates overfitting and improves generalization. At the same time, xNNPCD also outperforms the PCD-restricted baselines (“death”), indicating that known PCD gene sets alone are insufficient to explain cell fitness variation. The fully connected “others” neuron captures non-PCD signals that contribute to cell survival, complementing the pathway-specific channels.

Most importantly, xNNPCD shows a clear improvement over the static xNN, which uses the same pathway-constrained architecture without iterative mask refinement. This performance gain validates the iterative strategy: static binary annotations from public databases do not fully capture the complexity of gene-PCD relationships, and dynamically refining *M* from binary to continuous values allows the model to learn latent associations that improve predictive accuracy. This is further supported by the scatter plots in Supplementary Figure S1, where xNNPCD demonstrates a higher Pearson correlation coefficient (*r =* 0.554) than xNN (*r =* 0.541) between predicted and true dependency scores.

We noted that the absolute predictive performance (*R*^2^ ≈ 0.31) was moderate, which is expected given the noise and heterogeneity of the paired datasets. CMap L1000 assay directly measures only 978 landmark genes and infers the remaining 11,350, while Depmap dependency scores exhibit substantial variability across cell lines and sgRNA constructs. In addition, our sample-level split may place technical replicate signatures from the same gene-cell line pair in both training and test sets, which could modestly inflate absolute performance estimates. We therefore interpret the test-set *R*^2^ primarily as a comparative benchmark rather than a definitive estimate of out-of-sample performance for unseen knockouts or cell lines. Importantly, all baseline models were evaluated under the same split, and xNNPCD consistently outperformed fully connected and non-iterative alternatives, supporting the value of pathway-constrained architecture and iterative mask refinement.

### xNNPCD identifies candidate PCD regulators beyond prior annotations

The iterative training of xNNPCD converted the initial binary gene-PCD mask into a continuous-valued matrix. To evaluate whether this refinement preserves biological coherence, we compared learned mask values between prior-annotated and unannotated genes across all 12,328 CMap genes. Annotated genes showed significantly higher mask values than unannotated genes for all five PCD pathways (one-sided Wilcoxon rank-sum test, *p* < 0.0001; Figure 4). This pattern remained robust when the analysis was restricted to knocked-out genes with matched dependency scores, indicating that xNNPCD preserves curated pathway knowledge while extending it to additional candidate gene-PCD associations.

**Figure 4:**
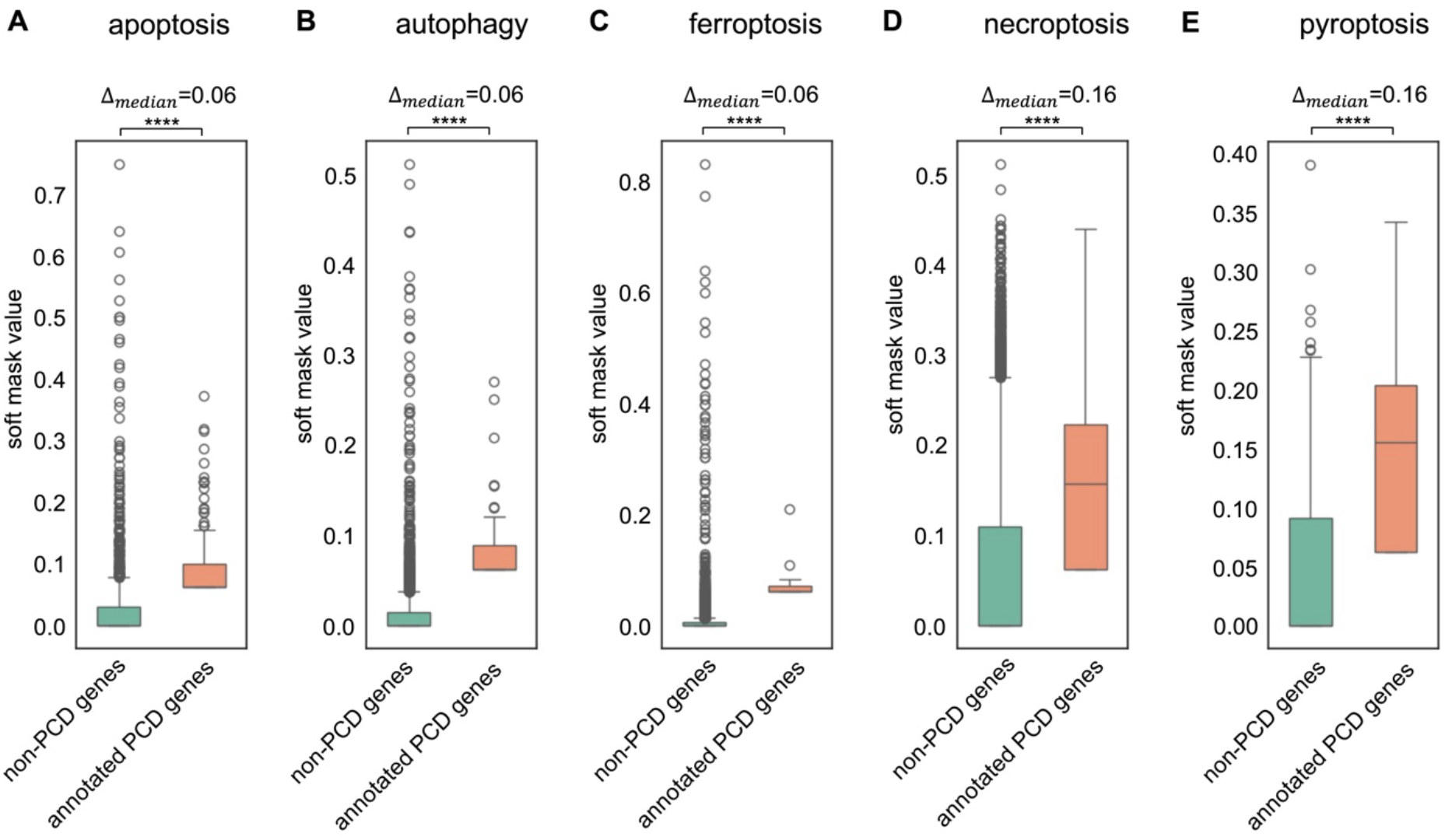
Learned soft mask values stratified by prior annotation for five PCD pathways. The boxplots illustrate the soft mask values learned by xNNPCD, stratified by prior annotation status. The distributions are shown for (**A**) apoptosis, (**B**) autophagy, (**C**) ferroptosis, (**D**) necroptosis, and (**E**) pyroptosis. Δ_*median*_ represents the difference in median values between the annotated and unannotated groups. Statistical significance was determined using a one-sided Wilcoxon rank-sum test, indicated by asterisks (****: *p* < 0.0001).

Notably, several genes without prior PCD annotations received high pathway-specific mask values, suggesting candidate gene-PCD associations not captured by current ontologies. Representative top-ranked unannotated candidates include *RPL23A* for apoptosis (rank 1, mask value 0.750), *HSPA5* for autophagy (rank 1, 0.511), *SNRPA1* for ferroptosis (rank 2, 0.775), *SLC6A2* for necroptosis (rank 2, 0.482), and *ASAH1* for pyroptosis (rank 1, 0.390). All five were perturbed in both CMap and DepMap, and each has independent experimental evidence supporting a connection to the corresponding pathway. He et al. (26) demonstrated that reduced *RPL23A* expression weakens its interaction with RPL11, thereby activating p53-mediated apoptosis, a mechanism distinct from its ribosomal function. Malek et al. (27) observed that *HSPA5* upregulation drives a protective autophagic response in drug-resistant myeloma cells. Similarly, Hao et al. (28) reported that the depletion of *SNRPA1* facilitates full p53 activation, which in turn induces ferroptosis in cancer cells. Regarding *SLC6A2*, studies indicate that its deficiency prevents norepinephrine reuptake, leading to an adrenergic surge that triggers RIPK1-RIPK3-dependent necroptosis (29,30). Finally, *ASAH1* functions as a negative regulator of pyroptosis by hydrolyzing ceramide, thereby preventing the lethal accumulation that drives ROS-dependent TXNIP/NLRP3 pathway activation (31,32). These examples support the ability of xNNPCD to nominate biologically plausible gene-pathway associations beyond standard annotations, while highlighting the need for targeted experimental validation to confirm direct regulatory mechanisms.

We next sought to understand why the model prioritized these specific candidates. Because the mask value reflects association with a specific death pathway, these genes are not nominated simply because they are essential; rather, they are prioritized when their knockout-induced transcriptional response is interpreted by xNNPCD as relevant to a particular PCD pathway. Dependency scores provide an additional layer of interpretation: candidates whose knockout strongly reduces fitness are more consistent with negative regulators, although essentiality alone cannot establish a direct PCD-suppressive role. Some genes, particularly ribosomal or housekeeping genes, may affect cell viability broadly and influence PCD-related programs indirectly.

### Mask values identify PCD regulators while essentiality distinguishes negatives from positives

Beyond nominating unannotated candidates, xNNPCD enhances the resolution of existing PCD annotations by replacing binary labels with continuous mask values. Analysis of the 3,967 knocked-out genes revealed that the mask value captures the strength of a gene’s functional coupling to a PCD pathway, not the direction of regulation: both negative regulators (which suppress death) and positive regulators (which promote death) may receive high mask values since they are tightly coupled to a pathway. The direction-specific biological outcome, whether losing a gene kills or spares the cell, is instead captured by the DepMap dependency score. These two metrics together support a two-step interpretive framework: mask values first identify genes with strong PCD pathway involvement, and dependency scores then discriminate negative from positive regulators within that group.

We first confirmed that the refined mask values correctly prioritize both classes of known regulators. As illustrated in Figure 5A, negative and positive PCD regulators both showed significantly higher mask values than non-PCD genes (one-sided Wilcoxon rank-sum tests: negative vs. non-PCD, *p =* 5.4 × 10^−9^; positive vs. non-PCD, *p =* 8.5 × 10^−7^), validating again that xNNPCD assigns strong pathway coupling to established regulators regardless of their functional direction. Notably, negative regulators showed slightly but significantly higher mask values than positive regulators (one-sided Wilcoxon rank-sum test, *p =* 0.0059). This partial directional signal arises naturally from the learning process: knockouts of negative regulators generate stronger and more directional fitness consequences, producing clearer ablation signals that the model attributes more consistently to specific PCD pathway neurons.

**Figure 5:**
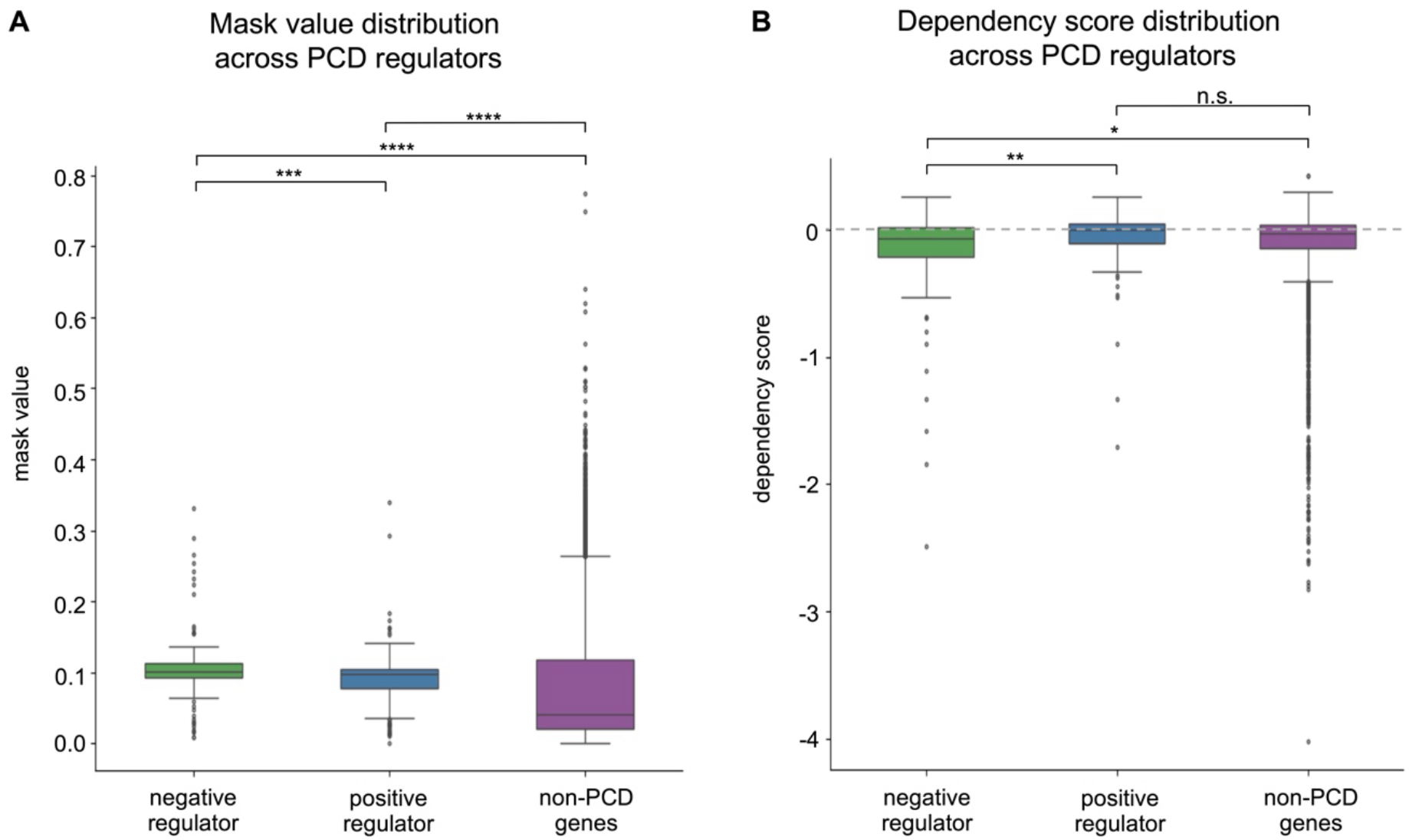
Comparative analysis of mask values and dependency scores across gene groups. (**A**) Mask value distributions for negative regulators, positive regulators, and non-PCD genes. (**B**) Dependency score distributions for the same groups. Statistical significance was determined using a one-sided Wilcoxon rank-sum test (n.s.: not significant, *: *p* < 0.05, **: *p* < 0.01, ***: *p* < 0.001, ****: *p* < 0.0001), except for the comparison between positive regulators and non-PCD genes in (**B**). The dashed line in (**B**) indicates a dependency score of 0.

We then applied dependency scores to complete the directional resolution. Negative regulators were significantly more essential than positive regulators, with markedly lower dependency scores (one-sided Wilcoxon rank-sum test, *p =* 0.0031, Figure 5B). Furthermore, negative regulators showed significantly lower dependency scores than non-PCD genes (*p =* 0.043), while positive regulators were indistinguishable from non-PCD genes (*p =* 0.64), confirming that essentiality is a specific signature of negative regulators rather than a general property of PCD pathway membership.

The complementary nature of the two metrics is further supported by their quantitative relationship. The Pearson correlation between mask values and dependency scores is −0.22 (*p* < 2.2 × 10^−16^) for annotated PCD genes, indicating that among pathway-annotated genes, stronger pathway coupling is associated with greater essentiality, while the correlation drops to −0.096 for non-PCD genes. This difference reflects the biological logic underlying the two-step framework: for PCD-annotated genes, high mask values preferentially mark negative regulators whose loss activates death, while no such structured relationship exists in the broader gene space. Together, a strong mask value coupled with a low dependency score forms a joint signature that specifically points to negative PCD regulators, a distinction that binary annotations alone cannot provide.

At the individual pathway level, significant essentiality stratification between high (top 25%) and low (bottom 75%) mask value groups was observed for apoptosis and autophagy (*p* < 0.0001, Supplementary Figure S2), but not for ferroptosis, necroptosis, or pyroptosis. This is biologically expected: apoptosis and autophagy are constitutively active under standard culture conditions, whereas the other three modalities require specific stress triggers absent from DepMap screens (5,33). The limited number of annotated knocked-out genes for ferroptosis (18 genes) and pyroptosis (15 genes) further reduces statistical power.

The global structure of the refined soft mask matrix is visualized in Figure 6, where a t-SNE projection of the five PCD pathway mask values for all knocked-out genes shows that apoptosis-related genes form a distinct cluster. This confirms that xNNPCD preserves the biological identity of well-characterized PCD pathways within the learned representation, complementing the regulator-class resolution demonstrated above.

**Figure 6:**
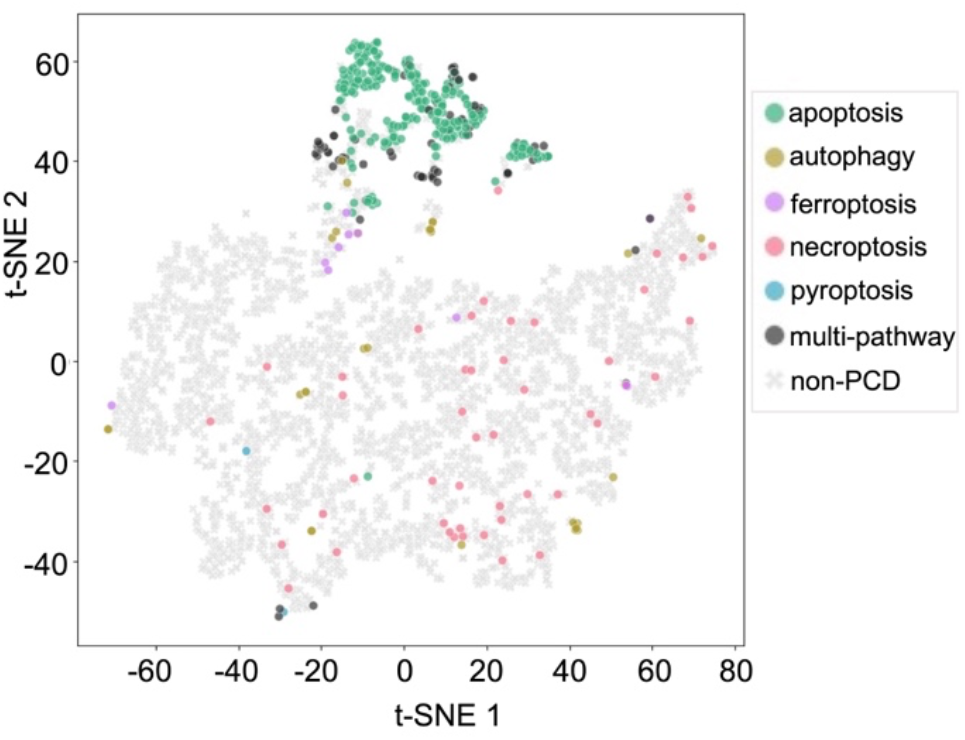
T-SNE projection of the learned soft mask matrix. Points represent knocked-out genes colored by prior pathway annotation. Dark gray dots indicate multi-pathway genes; light gray crosses indicate non-PCD genes.

### Transfer learning enables pathway-specific drug screening

To assess the translational utility of xNNPCD, we adapted the framework to predict CTRP drug sensitivity from CMap compound-induced differential expression signatures. We employed a transfer learning strategy in which both the refined mask matrix and the trained network weights from xNNPCD were used to initialize the drug sensitivity model.

As shown in Figure 7, training the model from scratch without transfer learning resulted in performance below a random baseline (*R*^2^ *=* −3.2^7^), meaning the model performed worse than simply using the mean drug sensitivity as the prediction for all samples. This confirms that the pharmacological dataset alone (6,665 samples) is insufficient to learn meaningful PCD-related structure *de novo*. Applying transfer learning drastically improved the predictive accuracy (*R*^2^ *=* 0.23, *MSE =* 0.014^7^), demonstrating that the PCD regulatory knowledge extracted from genetic perturbations transfers effectively to pharmacological contexts. While *R*^2^ *=* 0.23 remains modest in absolute terms, it represents a substantial improvement over the baseline and is consistent with the inherent difficulty of predicting drug sensitivity across heterogeneous cell lines from transcriptional profiles alone. More importantly, the objective of this analysis is not only global response prediction, but also pathway-level interpretation. In this setting, the refined mask matrix enables the model to assign drugs to biologically plausible PCD pathways with informative specificity.

**Figure 7:**
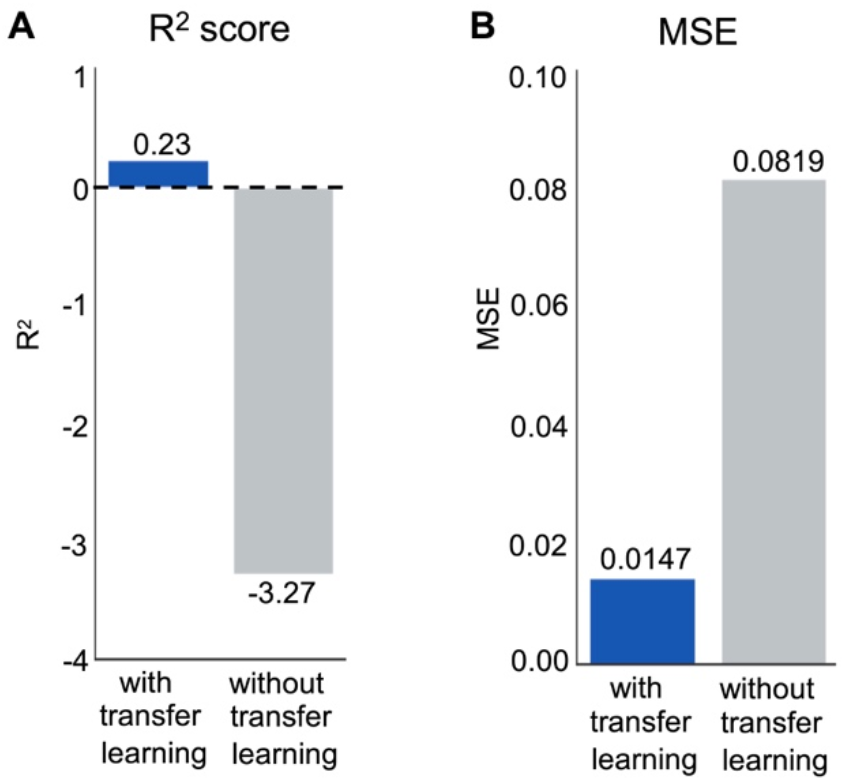
Performance comparison in predicting drug sensitivity. The bar plots compare the predictive performance of the model trained with and without the transfer learning strategy. The predictive performance is quantified by (**A**) the *R*^2^ score and (**B**) the Mean Squared Error (MSE). The dashed line in (**A**) indicates *R*^2^ being 0.

We next investigated pathway-level drug associations using the standardized activation scores (*Ã*_*d,p*_) across all evaluated drugs and PCD pathways (Figure 8). The highest-scoring drug-pathway pairs showed strong mechanistic plausibility. Drug decitabine emerged as the top candidate for ferroptosis (*Ã*_*d,p*_ *=* 10.39), consistent with evidence that decitabine induces reactive oxygen species accumulation, depletes glutathione, and suppresses GPX4 activity, thereby promoting ferroptosis in myelodysplastic syndrome cells (34). Compound BRD-K19103580 exhibited the highest overall activation score (*Ã*_*d,p*_ *=* 11.28), specifically targeting apoptosis. Although direct evidence linking this compound to apoptosis remains limited, BRD-K19103580 has been reported to exert additive anti-cancer effects on natural killer cells (35), which are known to induce target-cell apoptosis through Fas ligand and TRAIL pathways (36). These examples indicate that, even with modest sample-level predictive performance, the model can still recover biologically meaningful pathway assignments for individual compounds.

**Figure 8:**
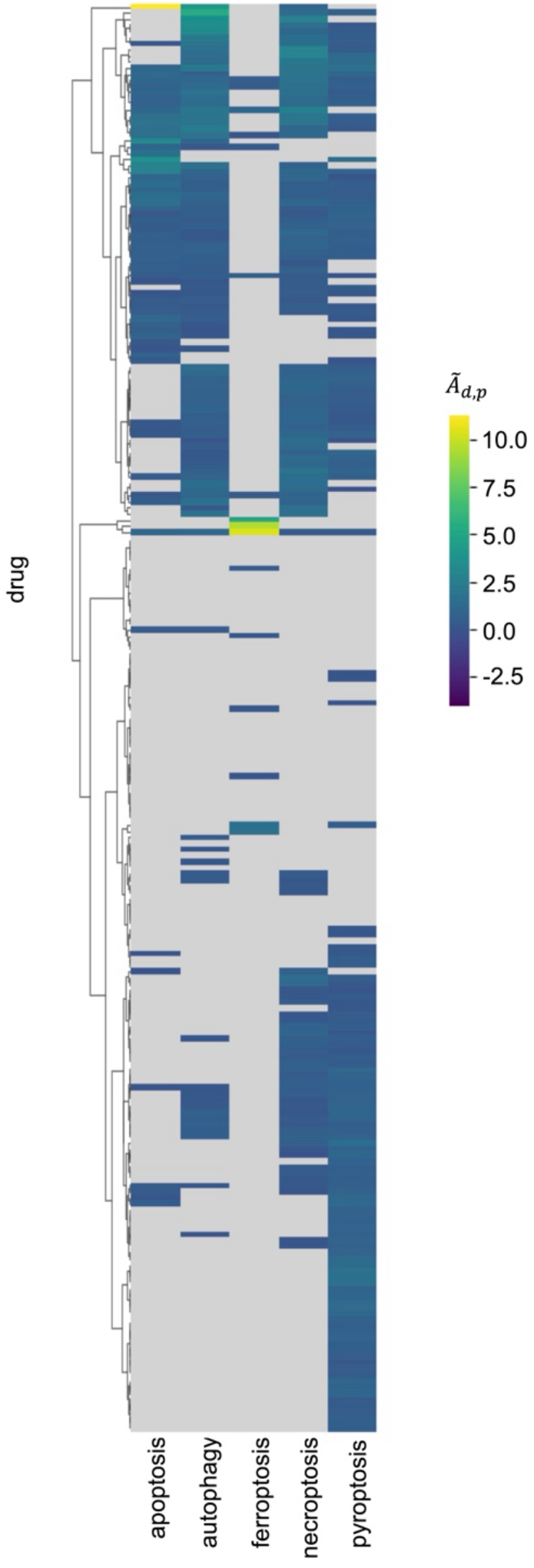
Clustered heatmap of drug-induced PCD pathway activation. The heatmap displays the standardized activation score (*Ã*_*d,p*_) for each drug across the five PCD pathways. Values where *Ã*_*d,p*_ < 0 are masked in gray to highlight above-average pathway activation. The dendrogram illustrates the hierarchical clustering of drugs based on their activation profiles.

Beyond prioritizing single agents, the pathway-resolved activation profiles also suggest a strategy for rational combination therapy design. Cancer cells frequently evade treatment by resisting one dominant death program, especially apoptosis (37). If xNNPCD can identify drugs that preferentially engage distinct PCD pathways, then combinations of agents targeting complementary death modalities may provide broader and more durable anti-cancer effects than single-pathway therapies alone. For example, pairing an apoptosis-biased compound with a ferroptosis-biased compound could simultaneously exploit multiple death vulnerabilities and reduce the chance of escape through pathway-specific resistance. Because Figure 8 reveals substantial diversity in pathway activation profiles across drugs, xNNPCD provides a principled basis for prioritizing candidate drug combinations for future experimental validation.

Together, these results show that the xNNPCD-derived mask matrix is useful not only for in silico identification of pathway-specific therapeutic agents, but also for guiding the rational design of PCD-targeted drug combinations.

## DISCUSSION

In this study, we developed xNNPCD as an interpretable framework for discovering regulators of programmed cell death from paired perturbation transcriptomic and fitness data. The central idea is to separate two related but distinct signals: transcriptional evidence that a gene is connected to a specific PCD pathway, and fitness evidence that loss of the gene compromises cell survival. This distinction allows xNNPCD to move beyond binary pathway annotations and clinical prognostic signatures toward pathway-level regulator discovery.

The refined gene-PCD matrix provides a quantitative view of pathway association. Unlike curated annotations, which usually indicate only whether a gene has been previously linked to a pathway, xNNPCD assigns continuous association scores between genes and individual PCD modalities. This enables known regulators to be ranked by pathway relevance and allows unannotated genes to emerge as candidate regulators. Importantly, the mask values should be interpreted as pathway-association scores rather than direct indicators of regulatory direction. Both positive and negative regulators can show high mask values if their perturbation affects a PCD-related transcriptional program. Directional interpretation requires the dependency phenotype: genes with high pathway association and strongly negative dependency scores are stronger candidates for negative regulators whose loss releases death signaling or otherwise compromises survival.

This separation between pathway association and regulatory direction is a major conceptual contribution of xNNPCD. It explains why the model can identify both positive and negative regulators from the refined mask, while preferentially prioritizing negative regulators through dependency scores. It also clarifies the biological meaning of the CMap-DepMap integration. CMap captures the molecular consequence of perturbing a gene, whereas DepMap captures the survival consequence of that perturbation. Neither readout alone is sufficient: expression signatures can suggest pathway involvement without indicating whether the perturbation is lethal, and dependency scores can identify essential genes without specifying the relevant death pathway. Their integration provides a more interpretable route to discovering PCD regulators.

The performance results should be viewed in this context. xNNPCD improved over fully connected and non-iterative baselines, supporting the value of imposing PCD pathway structure and refining it from data. However, the goal of xNNPCD is not simply to maximize prediction accuracy. The moderate predictive performance likely reflects the noise and heterogeneity of L1000 expression profiles and CRISPR dependency data, as well as the difficulty of predicting cell fitness from transcriptional responses alone. The value of the framework lies in using prediction as a scaffold for interpretable pathway-level inference.

The transfer to compound-induced perturbations illustrates a broader use of the learned representation. By assigning pathway-specific activation scores to drugs, xNNPCD can prioritize compounds that engage distinct PCD modalities. This is important because resistance to one death pathway, especially apoptosis, is a common feature of cancer. A pathway-resolved drug profile may therefore guide combination strategies in which compounds activating complementary PCD programs are paired to reduce pathway-specific escape. These predictions remain hypothesis-generating, but they provide a rational starting point for experimental drug-combination screens.

Several limitations should guide future work. First, xNNPCD can refine associations only for genes included in the CMap knockout dataset, so regulators outside this perturbation space remain inaccessible. Second, dependency scores are especially informative for negative regulators, whereas positive regulators may require gain-of-function perturbations, pathway activation assays, or context-specific stress conditions for clearer directional inference. Third, some high-scoring candidates may reflect broad essentiality rather than direct PCD regulation, requiring targeted perturbation and rescue experiments to establish the mechanism. Finally, future validation should test whether top-ranked genes and drug combinations produce pathway-specific cell-death phenotypes under appropriate biological contexts.

Overall, xNNPCD provides a framework for transforming large-scale perturbation data into interpretable maps of PCD regulation. By combining pathway-constrained modeling, iterative mask refinement, and dependency-based interpretation, it offers a route to discover both annotated and unannotated regulators and to extend PCD pathway analysis toward therapeutic prioritization.

## ACKNOWLEDGEMENTS

We thank all members of the Chen lab and Yingtong Liu for their discussions and suggestions throughout this study.

## AUTHOR CONTRIBUTIONS

Qingyang Yin: Conceptualization, Data curation, Formal analysis, Investigation, Methodology, Resources, Software, Validation, Visualization, Writing – original draft. Liang Chen: Conceptualization, Formal analysis, Funding acquisition, Investigation, Methodology, Project administration, Resources, Supervision, Writing – review & editing.

## SUPPLEMENTARY DATA

Supplementary Data are available at NAR online.

## DATA AVAILABILITY

The code is available at https://github.com/qyyin0516/xNNPCD. The processed datasets have been deposited in Zenodo and are accessible at https://doi.org/10.5281/zenodo.20060140. For raw data, the CMap differential expression signature data can be accessed from https://clue.io/data/CMap2020#LINCS2020. The DepMap dependency score data were downloaded from https://depmap.org/portal/download/all/ (version: 24Q2). The CTRP drug sensitivity data were downloaded from https://ctd2-data.nci.nih.gov/Public/Broad/CTRPv2.0_2015_ctd2_ExpandedDataset/.

