## Supplementary Data for "xNNPCD identifies regulators of programmed cell death by integrating perturbation transcriptomes with cancer dependency profiles"

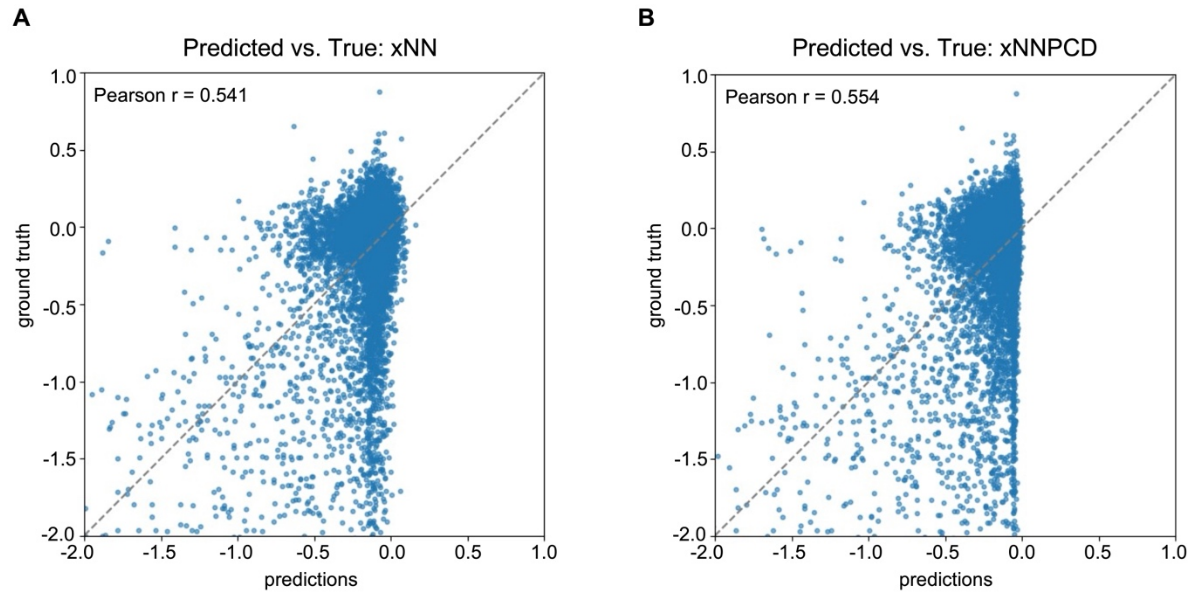


**Supplementary Figure S1. Scatter plots of predicted versus true dependency scores for xNN and xNNPCD.** The comparison is shown for (**A**) the static xNN and (**B**) the iterative xNNPCD. The dashed line represents the identity function $y = x$, indicating ideal prediction. The Pearson correlation coefficient ($r$) is displayed to quantify the alignment between predictions and ground truth.

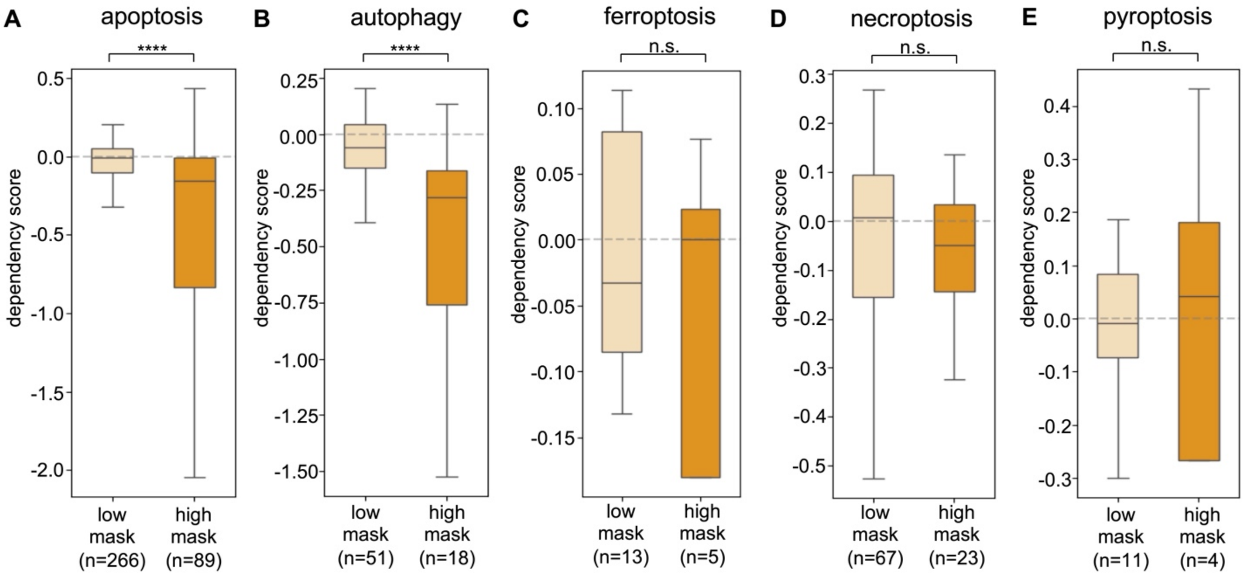


**Supplementary Figure S2. Stratification of dependency scores within individual PCD pathways based on learned mask values.** The boxplots illustrate comparisons of dependency scores between annotated knocked-out genes with high (top 25%) and low (bottom 75%) mask values for (**A**) apoptosis, (**B**) autophagy, (**C**) ferroptosis, (**D**) necroptosis, and (**E**) pyroptosis. Statistical significance was determined using a one-sided Wilcoxon rank-sum test (n.s.: not significant, ****: $p < 0.0001$). The dashed line indicates a dependency score of 0. Sample sizes ($n$) are indicated below each box.
